# Expression of the naked mole-rat transgene for *Has2* improved health span in C57Bl/6 mice, but it did not attenuate age-related hearing loss

**DOI:** 10.1101/2025.07.27.667071

**Authors:** Connor E. Owens, Aiden J. Borruso, Kevin A. Place, Andrei Seluanov, Vera Gorbunova, Patricia M. White, Ned J. Place

## Abstract

The naked mole-rat (NMR) is renowned for being the longest-lived rodent, with a maximum lifespan of over 40 years. This exceptional longevity has been attributed to increased cytoprotective signaling, which has been partially ascribed to a major component of the NMR’s extracellular matrix hyaluronan (HA), which has an unusually large molecular mass in this species. The HA molecular mass in the NMR is five to six times greater than in mice and humans. Mice expressing the transgene for NMR HA synthase 2 (*nmrHas2*), which is the enzyme that produces very high molecular mass HA (vHMM-HA), had improved health outcomes at an advanced age, such as improved mobility, grip strength, and reduced inflammation. Calorie restriction (CR) similarly improves healthspan in mice, and CR has also been shown to attenuate age-related hearing loss (AHL). Specifically, CR mitigates the effects of a single-nucleotide polymorphism (SNP) in the cadherin 23 (*Cdh23*) gene, which encodes a protein required for the stability of stereocilia on cochlear hair cells in mice and humans. Therefore, we hypothesized that AHL would be attenuated in *nmrHas2+* mice. We performed auditory brainstem response testing on 3-month and 12-month-old mice with (nmrHas2+) and without (nmrHas2-) the NMR transgene. We observed AHL at 12 months of age in both nmrHas2+ and nmrHas2-mice, and AHL was not attenuated by *nmrHas2* expression. This suggests that the cytoprotective effects of vHMM-HA work through a different pathway from CR, at least with respect to the inner ear.

## Introduction

The naked mole-rat (NMR) is a rodent renowned for its extreme longevity (40+ years) [1]. A mechanism that contributes to enhanced longevity in the NMR is increased cytoprotective signaling [2, 3]. This is partially attributed to the NMR’s very high molecular mass hyaluronan (also known as hyaluronic acid, vHMM-HA), which is an unbranched disaccharide glucuronic acid/N-acetylglucosamine polymer [2]. The NMR HMM-HA (6-12 MDa) is five to six times larger than mouse or human HMM-HA (0.5-3 MDa and 0.5-2 MDa, respectively) [2]. In mice and in humans, HMM-HA undergoes degradation and fragmentation, and when fragments are not properly cleared, this leads to the accumulation of low molecular mass (LMM) HA, which is biologically active and can induce inflammation and fibrosis [4-6]. The very high molecular mass HA (vHMM-HA) in NMRs is resistant to this degradation [7]. Additionally, binding of vHMM-HA to its receptor, CD44, induces expression of cytoprotective pathways and prevents binding of LMM-HA to CD44 [8].

A mouse line expressing the transgene for NMR hyaluronan synthase 2 (*nmrHas2*), which synthesizes vHMM-HA, was shown to have extended longevity, improved health outcomes and decreased inflamm-aging at 24 months of age, which was attributed to the reduced expression of inflammatory cytokines [9]. These results in the *nmrHas2* mouse are similar to the beneficial effects of calorie restriction (CR) [10]. The *nmrHas2* mouse is the subject of an ongoing project that aims to determine whether female reproductive aging is attenuated at 12 months of age after induction of the nmr*Has2* transgene at one month of age. As a supplement to the study on female reproductive aging, we added an investigation of age-related hearing loss (AHL), which is also evident by 12 months of age in the background strain (C57Bl/6) for the nmrHas2 mouse.

Age-related hearing loss, which can be alleviated in mice by CR, is typically caused by irreversible damage to cochleae that occurs during aging [11, 12]. In humans, approximately one in three individuals 65 to 74 years of age self-reports difficulty in hearing, including AHL [13]. Understanding potential mechanisms that contribute to or alleviate AHL is key to the development of successful therapeutics that attenuate AHL progression. This is increasingly important as global longevity increases, with average life expectancy increasing from 66.8 years in 2000 to 73.6 years in 2019 [14]. C57Bl/6 mice have pronounced AHL at 12 months of age, which is earlier than most other mouse strains, and is due to a single nucleotide polymorphism (SNP) in the gene cadherin 23 (*Cdh23*) [15]. Calorie restriction of C57Bl/6 mice at 12 months of age also prevented AHL [16].

Owing to the similar outcomes between *nmrHas2* expression and CR in healthspan and inflamm-aging, we anticipated *nmrHas2* mice would also demonstrate attenuated AHL. Our objective for this study was to perform ABR testing with *nmrHas2*+ and *nmrHas2*-mice at 3 and 12 months of age to test the hypothesis that AHL would be attenuated following induction of *nmrHas2* expression at 1 month. Following transgene induction, ubiquitous *nmrHas2* expression is maintained throughout life [9].

## Materials and Methods

### Animal husbandry

Mice were group housed and provided with standard chow diet and water *ad libitum*. All mice were housed under controlled conditions of temperature (range 21–24 °F), 12:12 h light:dark cycles, and relative humidity of 40 - 50%. All experimental procedures were approved by Cornell University’s Institutional Animal Care and Use Committee (protocol number 2020-0057).

### Transgenic animal design and use

Transgenic mice on the C57Bl/6 background were generated as previously described [9] (**Figure 1A**). We cross bred the *nmrHas2* and *R26*-*creER*^*t2*^ lines to generate heterozygous *nmrHas2* dams and homozygous *R26-creER*^*t2*^ sires. At 21 days of age, we weaned female offspring, placed an ear tag in one ear, and collected ear tissue from the opposite ear for genotype determination. Due to the primary use of these animals for female reproductive aging experiments, we euthanized male offspring at weaning.

**Figure 1:**
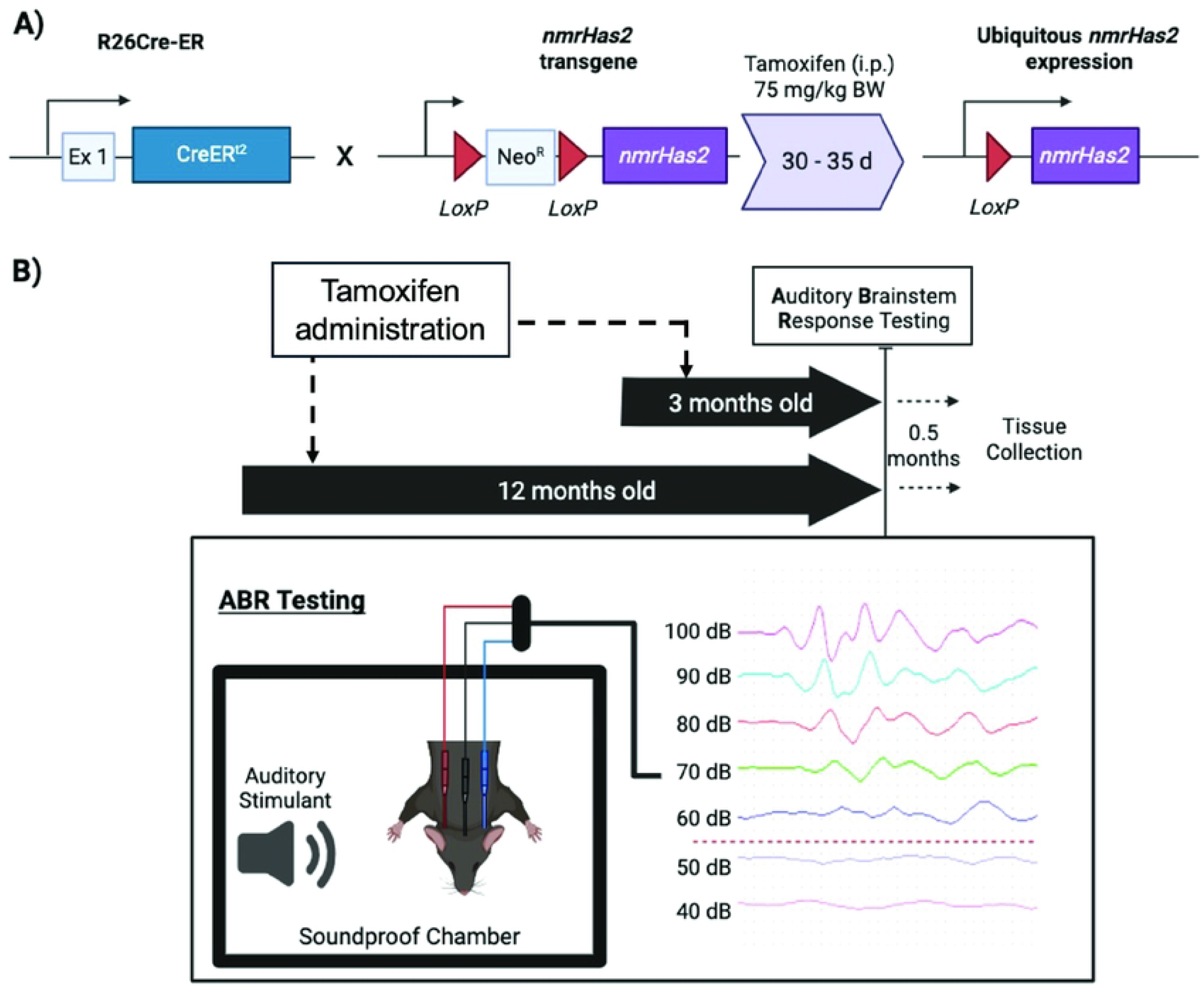
Outline of transgenic mouse model and experimental design. **A**) Experimental mice were generated by cross breeding sires that are homozygous for the CreER^t2^ recombinase in exon 1 of *Rosa26* (Cre-ER^t2^) with dams that are heterozygous for the *nmrHas2* transgene, for which the upstream Neo^R^-LoxP cassette prevents expression. Litters produced contained *nmrHas2+/-* x CreER^t2^+/- (nmrHas2+) and *nmrHas2-/-* x CreER^t2^+/- (nmrHas2-) offspring. One-month-old female offspring were given daily intraperitoneal injections of 75 mg/kg BW of tamoxifen for 5 days to induce *nmrHas2* expression in nmrHas2+ mice and equivalent doses of corn oil (Vehicle) to control for potential effects of tamoxifen. **B**) Mice were raised to 3 months or 12 months of age and underwent auditory brainstem response (ABR) threshold testing. The response threshold was determined as the lowest stimulus level (in decibels, dB) that elicited a wave-like pattern of response (in the example, it would be classified as 60 dB).

Experimental design is outlined in **Figure 1B**. We divided female mice into 2 groups based on genotype: *nmrHas2*+/-, *R26-creER*^*t2*^+/- (nmrHas2+, n = 13) and *nmrHas2*-/-, *R26-creER*^*t2*^+/- (nmrHas2-, n = 27). At 30 days of age, we administered 75 mg/kg tamoxifen (Tam; Sigma-Aldrich, St. Louis, MO) in sterile corn oil to mice via intraperitoneal (i.p.) injection daily for 5 days to ubiquitously induce life-time expression of *nmrHas2*. We administered equivalent doses of corn oil vehicle (Veh; Sigma-Aldrich, St. Louis, MO) to a subset of nmrHas2-mice to control for the potential effects of Tam treatment. Due to unanticipated expression of *nmrHas2* in ovaries of nmrHas2+ mice not administered tamoxifen, only nmrHas2+ mice given tamoxifen were included in these experiments. We raised mice to two target ages: 3 months (young, n = 15) and 12 months of age (old, n = 25). These timepoints correspond with previous literature for normal hearing and AHL in C57Bl/6 mice[12, 17]. Animals were separated into 6 groups based on genotype, age, and treatment combination: nmrHas2-× 3 m × Tam (n = 4), nmrHas2-× 3 m × Veh (n = 6), nmrHas2+ × 3 m × Tam (n = 5), nmrHas2-× 12 m × Tam (n = 9), nmrHas2-× 12 m ×Veh (n = 8), and nmrHas2+ × 12 m × Tam (n = 8).

### Auditory brainstem response (ABR) measurement and analysis

Approximately 2 weeks before reaching the target ages for euthanasia and tissue collection, we performed ABR testing on mice. Testing occurred between the hours of 9 AM and 12 PM EST. We deeply anaesthetized mice using an i.p. injection of ketamine (80 mg/kg animal weight) and xylazine (3 mg/kg animal weight). If after 10 minutes the desired plane of anesthesia had not been achieved (i.e. no response to foot pad pinch), an additional half dose of ketamine and xylazine was administered. We placed mice on a non-electric isothermal warming pad (Braintree Scientific, Inc., Braintree, MA) within a sound-proof chamber (Sound Room Solutions, Inc., Glen Cove, NY). We inserted three needle electrodes subcutaneously at the forehead and near each pinna. Electrodes were connected by cables to the ABR testing system (Tucker-Davis Technologies, Inc., Alachua, FL). We first evoked ABR responses using a 50 μsecond click stimuli. Immediately thereafter, we evoked ABR responses using 1 millisecond pure tones at frequencies of 4, 8, 12, 16, 24, and 32 kHz, with evoked responses averaged across 512 sweeps. We recorded responses for each stimulus level in 10 dB steps from 100 dB to 10 dB sound pressure level (SPL) for both clicks and tones. The ABR threshold was defined as the lowest SPL sufficient to elicit at least one peak that was consistent with an ABR waveform (**Figure 1B**). For the purpose of statistical analyses, we recorded animals that did not elicit a peak at any stimuli as having an ABR threshold of 110 dB, which is one 10 dB step above the maximum stimulus tested. A single researcher who was unaware of each animal’s age, Tam vs Veh treatment, and genotype scored ABR thresholds for all click and tone tests. Following the completion of the ABR testing and recovery from anesthesia, we returned mice to their pre-testing housing arrangement.

### Tissue collection, RNA extraction, and RT-PCR

Mice reached target ages approximately two weeks after ABR testing, at which time we individually euthanized mice via CO_2_ inhalation. Thereafter, we removed the cochleae by microdissection and processed them for future interrogation if the ABR results indicated AHL had been attenuated or alleviated. We also collected cochleae from a subset of young mice (n = 3 per genotype) for RNA extraction to confirm *nmrHas2* expression in the cochlea. For each mouse, we placed both isolated cochleae in a sterile 12-well plate on ice with chilled lysis buffer from the Qiagen RNeasy Plus Kit (QIAGEN; Hilden, Germany). We then broke apart cochleae using sterile forceps and scraped out sensory areas to expose cells to the lysis buffer. We further minced tissue first using sterile scalpel blades followed by homogenization in lysis buffer using Bead-Ruptor Pre-Filled Metal Bead Tubes (Omni International; Kennesaw, GA, USA). We extracted RNA from the homogenized tissue using the Qiagen RNeasy Plus Kit (QIAGEN; Hilden, Germany) and QIAShredder Columns (QIAGEN; Hilden Germany) according to manufacturer’s instructions.

We performed qRT-PCR to confirm cochlear expression of *nmrHas2* in nmrHas2+ mice. We used approximately 700 μg of extracted RNA to make cDNA using the SuperScript III First-Strand Synthesis kit (Invitrogen; Waltham, MA, USA). Due to the high similarity of *Has2* sequences in NMR and mice, we designed primers that were preferential to the NMR *Has2* mRNA sequence over the mouse *Has2* sequence, but not entirely exclusive. We performed qRT-PCR using primers preferential for *nmrHas2* (Forward primer: GAG-AGC-TCA-CAG-GTG-ACA-CAG; Reverse primer: GGA-GTC-ACA-AAC-CTG-CAC-ATA-ATC-CAC-A). We used previously generated cDNA libraries from NMR ovaries as a positive control for *nmrHas2* expression and primers specific for mouse *Has2* as a positive control for PCR in nmrHas2-mice. We performed gel electrophoresis and gel extraction using the Qiagen QIAQuick Gel extraction kit (Qiagen; Hilden, Germany) to isolate PCR products and confirm their sequences. We performed Sanger sequencing of PCR products at the Cornell Bioinformatics Resource Center using Big Dye Terminator Cycle (Applied Biosystems; Waltham, MA) and confirmed the sequence similarity of PCR products to the NMR *Has2* sequence.

### Statistical design

We performed all statical tests in JMP Pro, ver. 17.0.0 (SAS Institute Inc.; Cary, NC, USA). To assess if ABR threshold was different based on age, genotype, or tamoxifen treatment, we first performed a generalized linear model for clicks and all tones. The ABR threshold was the outcome variable and fixed effects were genotype, age, tamoxifen treatment, and genotype × age interaction. We performed the following planned post-hoc contrasts for clicks and all tones: 3 m and 12 m nmrHas2-, 3 m and 12 m nmrHas2+, and 12 m nmrHas2- and 12 m nmrHas2+. We did not observe an effect based on tamoxifen treatment, so this effect was not examined in post-hoc contrasts.

## Results

### ABR testing results

If AHL was attenuated by *nmrHas2* expression, this would manifest as decreased ABR thresholds. For clicks and all tones, ABR thresholds were significantly different based on age (*P ≤* 0.004, **Figure 2, Table S1**), but not based on genotype (*P ≥* 0.37). Planned post-hoc contrasts demonstrated the mean ABR thresholds were greater in old nmrHas2-mice for clicks and all tones compared to young nmrHas2-mice (*P ≤* 0.006, Table S1). The ABR threshold was greater in old nmrHas2+ mice for clicks and tones at 4, 8, 12, 16, and 24 kHz compared to young nmrHas2+ mice (*P ≤* 0.001), but did not achieve statistical significance at 32 kHz (*P* = 0.08, Table S1). The ABR threshold was also not different between old nmrHas2- and nmrHas2+ mice for clicks (*P* = 0.26, **Figure 2A**) and all tones (*P ≥* 0.26, **Figure 2B**).

**Figure 2:**
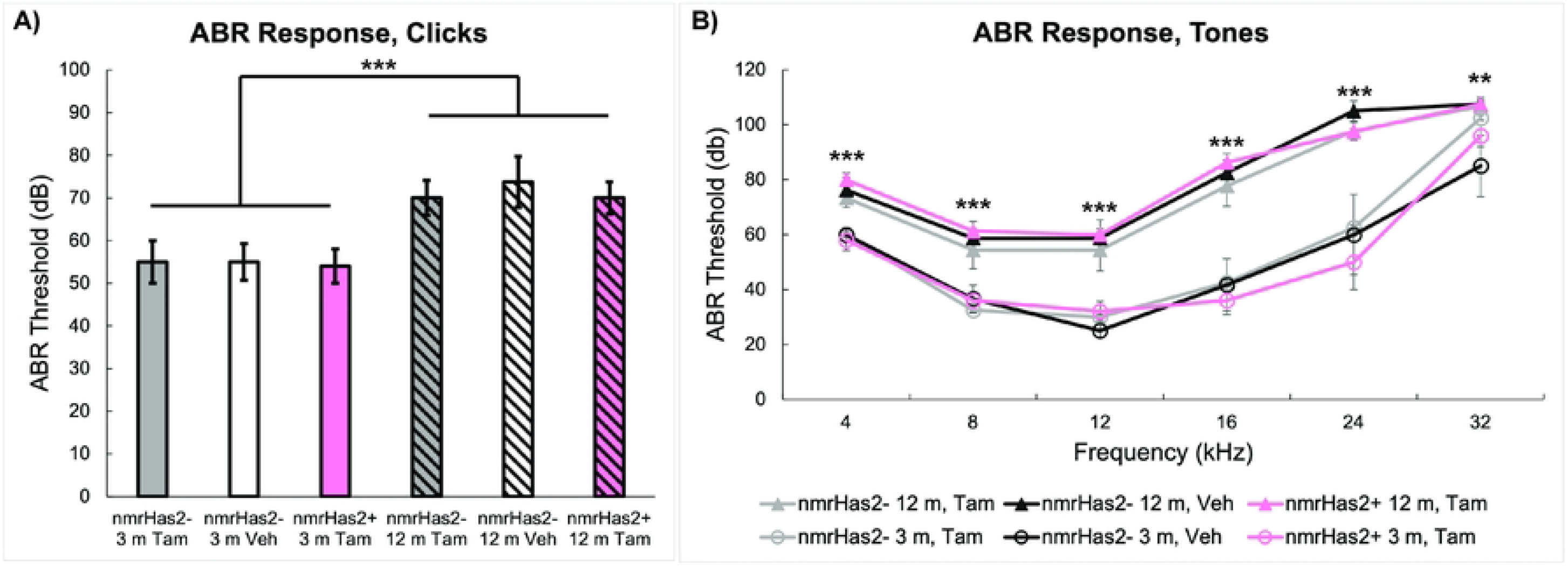
Results of auditory brainstem response (ABR) testing when 3- and 12-month-old mice that were stimulated with **A)** clicks and **B)** pure tones. Expression of naked mole-rat hyaluronan synthase 2 was induced in mice carrying the transgene (nmrHas2+) using intraperitoneal tamoxifen (Tam) injections at 1 month of age. Control mice without the transgene (nmrHas2-) were given either Tam or corn oil vehicle (Veh). The ABR threshold was higher for clicks and tones in old mice compared to young mice, regardless of genotype or tamoxifen treatment, indicating *nmrHas2* did not attenuate age-related hearing loss. For differences based on age, ^**^ indicates *P <* 0.01 and ^***^ indicates *P* < 0.001.

### Cochlear expression of nmrHas2

We used extracted RNA to validate expression of *nmrHas2* in nmrHas2+ mice. Zhang et al. (2023) established overall *Has2* tissue expression was substantially greater in *nmrHas2+* mice following Tam induction^22^. Therefore, we used qRT-PCR to evaluate cochlear *Has2* expression. Primers preferential for *nmrHas2* yielded a single-band product from NMR ovarian cDNA libraries while *musHas2* primers yielded no product from the same NMR libararies. Expression of *Has2* was substantially greater in the cochleae of nmrHas2+ mice compared to nmrHas2-mice (Fold change = 781.98 ± 234.89 and 1.00 ± 0.38, respectively). These data indicated successful induction of expression of *nmrHas2* in the cochleae of nmrHas2+ mice.

## Discussion

The goal of this experiment was to determine whether ubiquitous expression of *nmrHas2* transgene, including within the cochleae, attenuated AHL. Ubiquitous expression of *nmrHas2* in mice attenuated similar age-associated phenotypes to those previously seen using CR, such as improved overall health and reduced inflammation [9] Calorie restriction has also been demonstrated to attenuate AHL in mice at 12 months of age [16]. Therefore, we hypothesized that ubiquitous expression of *nmrHas2*, which includes cochleae, would attenuate AHL at 12 months of age in mice. While we demonstrated cochlear expression of the *nmrHas2* transgene in nmrHas2+ animals, there was no attenuation of AHL in nmrHas2+ mice at 12 months of age compared to nmrHas2-mice.

While the specific cytoprotective mechanisms induced by vHMM-HA in the NMR are not well understood, the cytoprotective pathway regulated by the transcription factor nuclear erythroid 2-related factor 2 (NRF2) is upregulated in the NMR relative to mice [3]. Typically, Kelch-like ECH-associated protein 1 (KEAP1) sequesters NRF2 in the cytoplasm and leads to NRF2 ubiquitination and degradation [18]. However, during periods of stress, a conformational change in KEAP1 allows NRF2 translocation to the nucleus to upregulated cytoprotective pathways [19]. In the NMR, KEAP1 is downregulated, and NRF2 and its downstream targets are continuously upregulated [3]. In a KEAP1 knock down mouse model, the NRF2-regulated pathways were continuously upregulated, and AHL in C57Bl/6 mice was attenuated at 12 months of age [20]. Conversely, loss of NRF2 in mice accelerated progression of AHL due to earlier loss of cochlear hair cells [21]. As a caveat, the upregulation of NRF2 in the cochlea from vHMM-HA expression remains to be determined. Nonetheless, these studies suggested that the *nmrHas2* transgene would mitigate AHL in aging mice.

As this short report demonstrates that *nmrHas2* is not sufficient to overcome AHL in aging C57BL/6 mice, we may speculate on the underlying mechanisms. First, *nmrHas2* generates vHMM-HA, which binds to the widely expressed CD44 receptor [8]. In the cochlear sensory epithelium, only outer pillar cells express CD44 [22]. Previous reports indicate that CD44 is necessary for the anti-aging effects of vHMM-HA [8]. Our data indicate that cochlear hair cells were not rescued by the changes in the extracellular environment, possibly because they do not express CD44. Alternatively, NRF2 also plays key roles in cholesterol metabolism [23]. Cholesterol is a necessary component of the outer hair cell amplification mechanism involving SLC26A5 [24]. Loss of outer hair cell amplification through changes in cholesterol metabolism could potentially reverse any benefits derived from NRF2 activation or reductions in inflammation.

The early onset of AHL in C57Bl/6 mice enabled us to add ABR testing to our larger study on the effects of the *nmrHas2* transgene on female reproductive aging at 12 months of age. It appears that, unlike CR, the anti-inflammatory properties of vHMM-HA observed in previous studies using nmrHas2 mice [9] could not overcome the hair cell intrinsic hearing loss caused by the variant SNP in *Cdh23*. One potential reason may be that while *nmrHas2* was expressed in the cochlea, it did not generate sufficient vHMM-HA to attenuate AHL.

For future studies, a different mouse strain in which AHL occurs through a different mechanism could be used. For example, AHL in CD/1 mice has been attributed to inflammation-induced aging of the cochleae [25]. Therefore, CD1 mice might be informative in accessing the potential cytoprotective effects of *nmrHas2* expression in the cochlea. As inflammation is a common sequelae to noise or drug damage, it will be interesting to see if vHMM-HA can mitigate hearing loss in damage paradigms through its action on endogenous cochlear macrophages [26]. Lastly, it would be interesting to see if vHMM-HA can alleviate the oxidative stress observed in the mouse cochlea from exogenous factors, such as cigarette smoke exposure [27].

Overall, we did not observe an attenuation of AHL in mice that ubiquitously express *nmrHas2*. However, these data raise questions regarding the mechanism of the cytoprotective effect of vHMM-HA, as only outer pillar cells, and distinctly not outer hair cells, express CD44, a primary receptor for HA. Future studies examining cochlear cell-type specific HA-interactions and exposure to various exogenous factors that cause hearing loss could elucidate the primary cyptoprotective mechanism associated with *nmrHas2* expression.

## Acknowledgements

We would like to thank our lab technicians, Rebecca Cubitt and Ruby Feng, for their help in maintaining the *nmrHas2* and *Cre-ER*^*T2*^ mouse colonies and i.p. injections to induce expression of the transgene. We would also like to thank our undergraduate researchers, Kristen Dookie, Andy Zhang, Shruti Nagpal, and Posie Price, for their help maintaining the mouse colonies and assisting during tissue collection.

